# Punctuated evolution shaped modern vertebrate diversity

**DOI:** 10.1101/151175

**Authors:** Michael J. Landis, Joshua G. Schraiber

## Abstract

The relative importance of different modes of evolution in shaping phenotypic diversity remains a hotly debated question. Fossil data suggest that stasis may be a common mode of evolution, while modern data suggest very fast rates of evolution. One way to reconcile these observations is to imagine that evolution is punctuated, rather than gradual, on geological time scales. To test this hypothesis, we developed a novel maximum likelihood framework for fitting Lévy processes to comparative morphological data. This class of stochastic processes includes both a gradual and punctuated component. We found that a plurality of modern vertebrate clades examined are best fit by punctuated processes over models of gradual change, gradual stasis, and adaptive radiation. When we compare our results to theoretical expectations of the rate and speed of regime shifts for models that detail fitness landscape dynamics, we find that our quantitative results are broadly compatible with both microevolutionary models and with observations from the fossil record.

---

A key debate in evolutionary biology centers around the seeming contradictions regarding the tempo and mode of evolution as seen in fossil data compared to ecological data. Fossil data often support models of stasis, in which little evolutionary change is seen within lineages over long timescales (Hunt 2007; Estes and Arnold 2007), while ecological data show that rapid bursts of evolution are not only possible, but potentially common (Reznick et al. 1997; Herrel et al. 2008; Brown and Brown 2013; Cresko et al. 2004). At face value, these observations seem to contradict one another, an observation known as the “paradox of stasis” (Hansen and Houle 2004). These observations are often reconciled through a descriptive model of punctuated evolution, entailing stasis interrupted by pulses of rapid change, as famously articulated by Simpson (Simpson 1944; 1953).

On macroevolutionary timescales, pulses of rapid change are expected to look roughly instantaneous. Only recently have statistical methods grown sophisticated enough to model punctuated evolution as a stochastic process, with advances showing that punctuation is detectable in some fossil time series (Hunt et al. 2015) and between pairs of living and extinct taxa (Uyeda et al. 2011).

While these studies establish the existence of punctuated evolution, it is still unknown whether the evolutionary mode is common or rare. How many clades in the Tree of Life were shaped by abrupt pulses of rapid evolution? If these evolutionary pulses are common, then that should inform our expectations about how traits evolved for clades that left no fossils, and the potential for vulnerable species to adapt rapidly to climate change (Quintero and Wiens 2013). To this end, phylogenetic models—models of trait evolution that account for the shared ancestry of species—have played a vital role in measuring the relative support of competing Simpsonian modes of evolution.

A pioneering meta-analysis (Harmon et al. 2010a) fitted a collection of phylogenetic models to 49 animal clades, finding preference for gradual and stationary modes of evolution. Their work predated the advent of phylogenetic models of punctuated evolution (Landis et al. 2013; Duchen et al. 2017b), so its frequency could not have been measured. Moreover, there remains a concern that punctuated and gradual models leave similar patterns of trait variation in neontological data (Hansen and Martins 1996; Clavel and Morlon 2017). While recent methodological developments show that there is power in comparative data to identify punctuated evolution (Bokma 2002; Landis et al. 2013; Duchen et al. 2017a), little is known about either the statistical properties of these methods, nor is much known about the prevalence of punctuated change throughout some of Earth’s most intensely studied clades.

Here, we examine evidence for punctuated evolution across vertebrate taxa using a method for fitting Lévy processes to comparative data. These processes can capture both gradual and punctuated modes of evolution in a single, simple framework. We apply this method to analyze 66 vertebrate clades containing 8,323 extant species for evidence of punctuated evolution by comparing the statistical fit of several varieties of Lévy jump processes (modeling different types of punctuated evolution) to three models that emphasize alternative macroevolutionary dynamics. Under these models, the adaptive optimum of a lineage may wander gradually and freely (Brownian motion), it may change gradually but remain stationary (Ornstein-Uhlenbeck), or it may change most rapidly following the initial diversification of a clade while decelerating over time, e.g. during an adaptive radiation (Early Burst). Beyond simple model comparison, we show that the parameter estimates corresponding to the microevolutionary and macroevolutionary mechanisms of the model have biologically meaningful interpretations (Estes and Arnold 2007), illuminating previously hidden features of Simpson’s adaptive grid.

## 1 Results

### 1.1 Maximum likelihood method has power to distinguish punctuated evolution from comparative data

We developed a maximum likelihood method for fitting Lévy processes to phylogenetic comparative data using restricted maximum likelihood estimation (REML), by analyzing the phylogenetically independent contrasts (Felsenstein 2004) (*Materials and Methods*). The Lévy processes we apply in this work consist of two components: a Brownian motion and a pure jump process. The Brownian motion is characterized by a rate parameter, *σ*^2^, and the pure jump process is characterized by a Lévy measure, *v*(*·*), where *v(dx)dt* can be thought of as the probability of a jump with a size in the interval (*x*, *x* + *dx*) occuring in the short time *dt.*

Both Lévy processes with jumps and pure Brownian motion accumulate variance proportional to time (*SI Appendix*), leading to speculation that it is impossible to distinguish between punctuated and certain gradual models from comparative data (Hansen and Martins 1996; Clavel and Morlon 2017). For simulations with moderately sized clades (>100 taxa), we had sufficient power to differentiate punctuated evolution from other Simpsonian modes of evolution. This is possible due to the impact of rare, large jumps that cause a heavy-tailed distribution of trait change (*SI Appendix*). Moreover, we saw low false positive rates for identifying punctuated evolution, even in the presence of phylogenetic error (4% for clades with ~100 taxa, 7% for clades with ~300 taxa; *SI Appendix).*

### 1.2 Extant vertebrate body sizes evolved by rapid bursts

We assembled comparative datasets from time-scaled tree estimates and body size measurements for 66 clades across five major vertebrate groups (Table 1). We computed Akiake Information Criteria weights (wAIC) (Akaike 1981) for each dataset from a panel of models. We used a Brownian motion (BM) to model gradual phenotypic evolution, in which a lineage’s phenotype follows a wandering optimum. We used the Ornstein-Uhlenbeck (OU) model to gradual evolutionary stasis around a single optimum and the Early Burst (EB) model to capture adaptive radiation, in which the tempo of evolution slows over time. We also compared two different types of jump processes: the compound Poisson with normally distributed jumps (JN or Jump Normal) and the Normal Inverse Gaussian (NIG). The JN process waits for an exponentially distributed amount of time before jumping to a new value; we use this to represent stasis followed by large-scale shifts between adaptive zones. On the other hand, the NIG process is constantly jumping, with larger jumps requiring longer waiting times; this model captures the dynamics of constant phenotypic change, where the majority of change occurs within the width of an adaptive zone, but rare shifts between adaptive zones are still possible. Finally, we examined the combination of the Brownian and jump processes, BM+JN and BM+NIG. All body size measurements were affected by intraspecific noise, which we fitted with the normal distribution. Examples of how each different Lévy process models trait evolution can be found in the *SI Appendix.*

**Table 1:** Vertebrate datasets. Data were collected for *N* = 66 clades for 8323 species records in total (mean 126 species per clade). The “Phylogeny” column indicates how phylogenies within each vertebrate group were generally acquired: by pruning from a large tree or by collecting independently estimated trees from the literature.

|  | # clades | Clade sizes |  | Phylogeny | Body size |
| --- | --- | --- | --- | --- | --- |
|  |  | mean | (min, max) |  |  |
| Fish | 12 | 105 | (37, 251) | Collected (Harmon et al. 2010b; Cowman and Bellwood 2011; Santini et al. 2013a;b; Sorenson et al. 2013; Bloom and Lovejoy 2014; Roxo et al. 2014; Campanella et al. 2015; Dornburg et al. 2015; Martin and Bonett 2015) | Length from (Froese and Pauly 2000) |
| Amphibians | 4 | 176 | (103, 231) | Pruned from (Pyron and Wiens 2011; Pennell et al. 2014) | Length from (De Lisle and Rowe 2013) |
| Reptiles | 24 | 127 | (53, 220) | Pruned from (Wright et al. 2015; Jaffe et al. 2011) | Length from (Feldman et al. 2016) |
| Birds | 17 | 111 | (56, 260) | Pruned from (Jetz et al. 2012) | Mass from (Dunning Jr 2007) |
| Mammals | 9 | 158 | (69, 229) | Collected (Vos and Mooers 2004; Slater et al. 2010; Nyakatura and Bininda-Emonds 2012; Bibi 2013; Mitchell et al. 2014; Schenk et al. 2013; Shi and Rabosky 2015) | Mass from (Jones et al. 2009) |
| Total | 66 | 126 | (37, 260) | – | – |

Figure 1 shows that that 32% of clades were best fit by models of punctuated evolution. Brownian motion, Early Burst, and Ornstein-Uhlenbeck were each selected substantially less often (18%, 15%, and 2%, respectively). 33% of clades showed little support for any single mode of trait evolution. We did not find any evidence that a particular evolutionary mode is enriched in any of the five vertebrate groups (chi-squared test, p=0.08). Lévy jump models and Brownian models fitted to any given clade yield nearly identical estimates of process variance (*SI Appendix*), indicating that the tempo and mode of evolution leave distinct patterns in comparative data.

**Figure 1:**
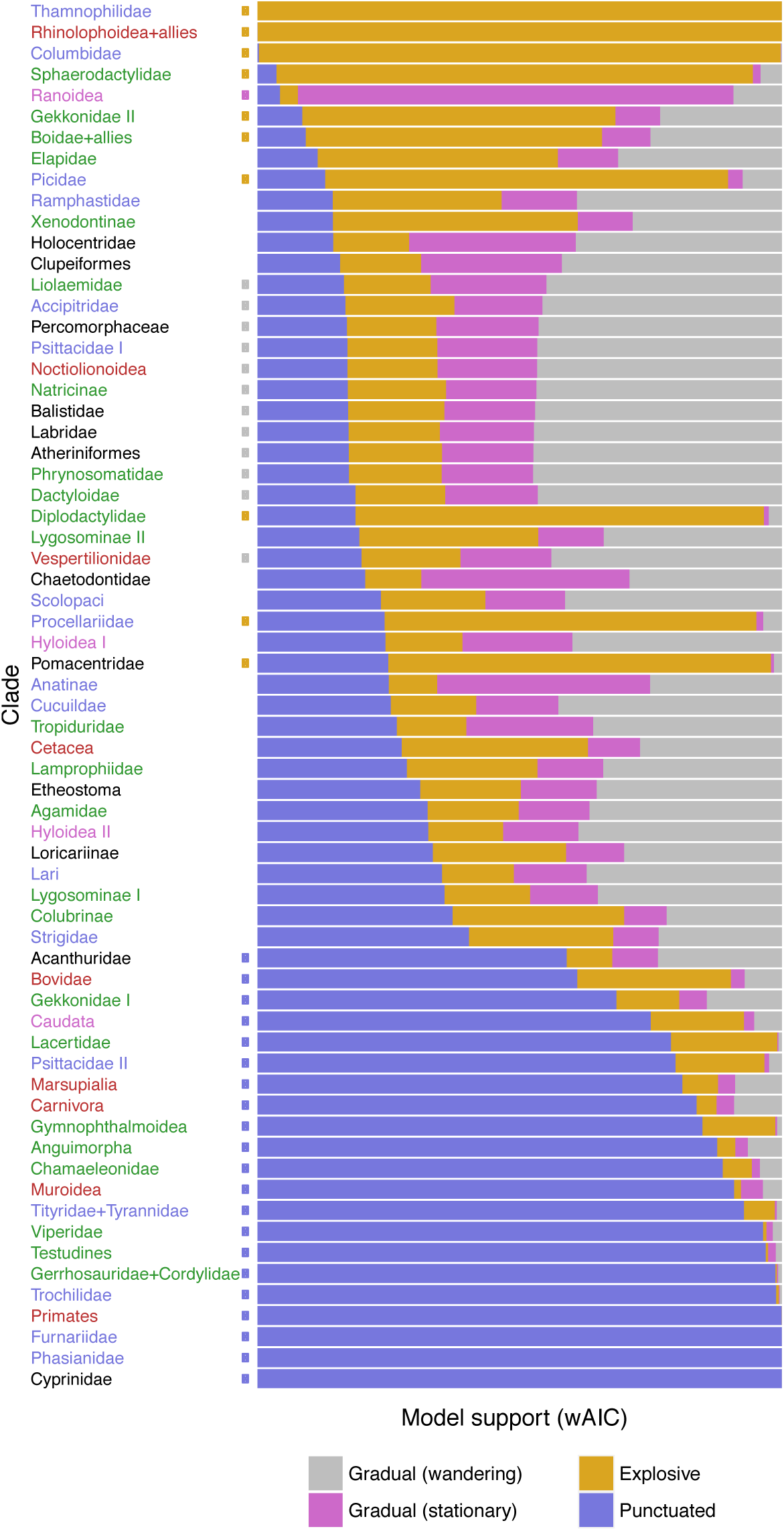
Model selection profiles for 66 vertebrate clades. Clade colors indicate their order: black, fish; purple, amphibians; green, reptiles; blue, birds; red, mammals). Each clade was fit to seven models, classified into four groups: Gradual-wandering (BM), Gradual-stationary (OU), Explosive (EB), and Punctuated (JN, NIG, BM+JN, BM+NIG). AIC weights were computed using only the best fitting model within each class. A model class is selected only if its AIC weight is twice as large than that of any other model class (circles indicate selection counts: 12 Gradual (wandering), 1 Gradual (stationary), 10 Explosive, 21 Punctuated, 22 ambiguous). Alternative model classifications are provided in *SI Appendix.*

We speculated that clades would favor models that produced similar patterns in comparative data. To characterize this, we applied a principal components analysis to the wAIC scores of Figure 1, then clustered wAIC profiles using k-means (k=4). The first three principal components explain 85.8% of the variance. Figure 2 shows that four clusters form around clades that select models of gradual evolution (Brownian motion and Ornstein-Uhlenbeck), explosive evolution (Early Burst), and the two models representing punctuated evolution (Jump Normal and Normal Inverse Gaussian). Interestingly, the third component separates clades that favored the two different kinds of jump models we explored, with the most punctuated process (Jump Normal) running opposite to the infinitely active processes (Normal Inverse Gaussian). This suggests that with more power it will be possible to choose among different models of punctuated evolution with greater confidence.

**Figure 2:**
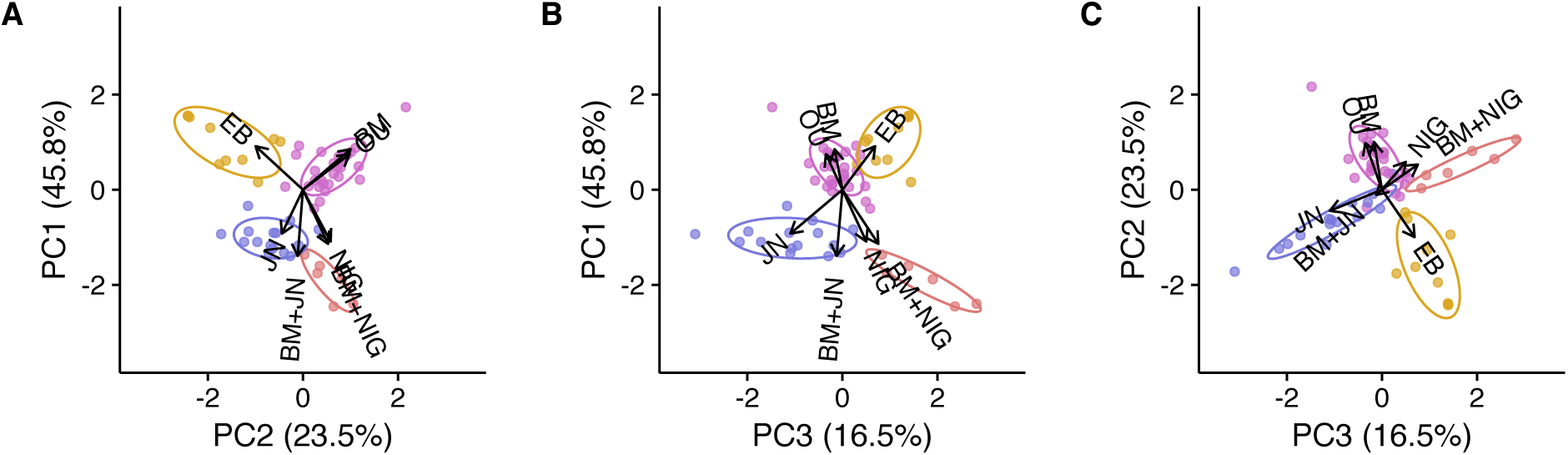
Empirical axes of trait evolution. The principal components and k-means clusters (*k* = 4) for wAIC scores reported in Figure 1. Panels A, B, and C plot the first three principal component axes as pairs, explaining 85.8% of the model-fit variance. Both PCA and the k-means clusters identify three major axes of trait evolution: gradual-wandering and gradual-stationary (purple), explosive (gold), and punctuated models, with the punctuated axis being divisible into subtypes of jump processes with finite (blue) and infinite (red) activity.

### 1.3 Waiting times to transition between adaptive zones

Under a microevolutionary model of stabilizing selection, intraspecific variation is distributed normally around the adaptive optimum (Lande 1976). This intraspecific variation in turn approximates the width of a given adaptive zone (Simpson 1944; Van Valen 1973). When fitting models of trait evolution, we inferred a parameter that corresponds to the standard deviation of intraspecific variation. Because we have only one sample per species, this is a noisy measure of intraspecific variation that also includes measurement error. We therefore regressed our estimates of the intraspecific variation using bird data with known phenotypic standard deviations, and used the regressed estimates as a measure of intraspecific variation, *σ*_intra_ (see Methods).

We sought to understand the expected waiting time until lineages escape their current adaptive zones. To do so, we computed the expected waiting time, 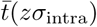, until the lineage jumps at least *z* intraspecific standard deviations away from the current phenotype (*Materials and Methods*). Because the Jump Normal tends to be slightly preferred among jump models, we analyzed all clades using Jump Normal (Figure 3A). The value at *z* = 0 shows the waiting time for a jump to occur at all, while we take *z* > 2 to be a jump outside of the current adaptive zone, which we define as the interval containing roughly 95% of phenotypic variation per lineage. We found that lineages wait between 1 and 100 million years before shifting to new adaptive zones, with most lineages waiting on the order of 10 million years between shifts (median waiting time 10^7.4^ years for *z* = 2).

**Figure 3:**
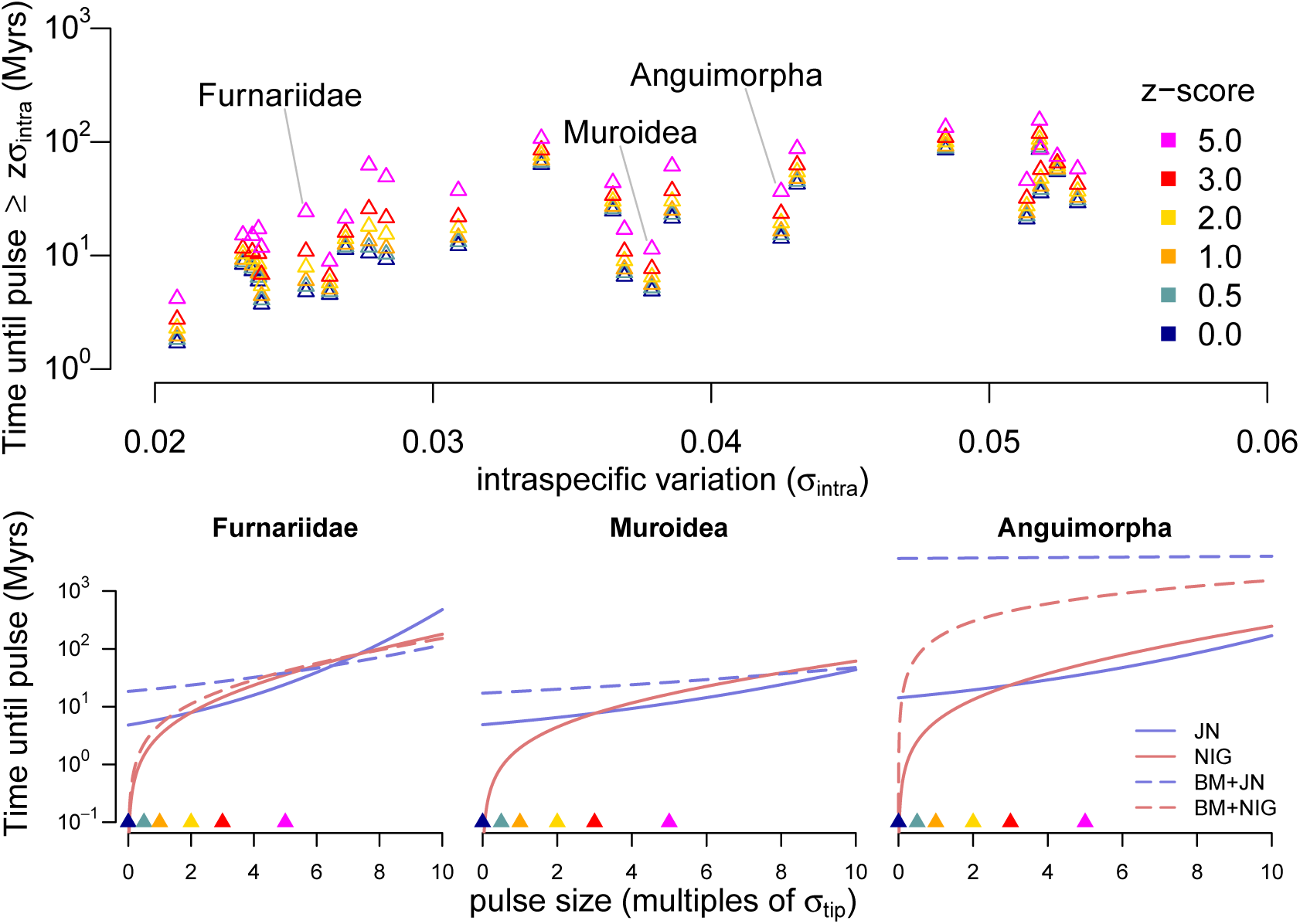
Expected waiting time to shift between adaptive zones. A) Plot shows the expected waiting of time needed to produce a evolutionary pulse that is at least *z* times that of the standard deviation of the intraspecific variance (*σ*_intra_). Times shown for the pure jump-normal process for the 25 clades that select jump processes and *σ*_intra_ > 0 by any amount. B) Waiting times until pulses at least as large as given size differs between the four fitted jump models for three clades shown in (A). Jump-normal (blue) has finite activity, and cannot easily produce small amounts of change quickly, unlike the normal-inverse Gaussian (red), which is infinitely active. AIC selects pure jump models (solid lines) for *Furnaridae* (left) and *Muroidea* (center), and a jump-diffusion model (dotted lines) for *Anguimorpha* (right).

The waiting time between adaptive shifts depends both on the magnitude of the shift and on the underlying jump model (Figure 3B). Models with jumps differ in two ways: 1) the relative frequencies of jumps within and between adaptive zones (e.g. *z* < 2 vs *z* ≥ 2), and 2) whether the fitted model is a pure jump process or a jump-diffusion process. JN models experience long periods of stasis, and jumps tend to be large. Thus, small changes within a lineage’s adaptive zone occur on approximately the same time scale as jumps between adaptive zones. In contrast, infinitely active processes, as represented by NIG, jump continuously. Under these models, shifts within an adaptive zone occur frequently, which could reflect rapid adaptation to small-scale environmental perturbations, while larger shifts between adaptive zones occur on much longer time scales. Interestingly, we find that both finite and infinite activity pure jump models predict similar rates of shifting between adaptive zones, as indicated by the fact that the solid curves are near each other for *z* ≥ 2. When looking at a clade that is best fit by a jump-diffusion model, such as *Anguimorpha* (Figure 3B, right), we find that they explain most of the morphological disparity among taxa via the Brownian motion component of the model, and we see that jumps occur extremely rarely compared to pure jump models. Jump-diffusion models with small diffusion components produce jump size-time curves that are essentially identical to the jump size-time curves for the corresponding pure jump model (compare NIG and BM+NIG for *Muroidea*).

## 2 Discussion

The pulses of punctuated evolution proposed by Simpson (Simpson 1944; 1953) are evident in ecological (Thompson 1998; Carroll et al. 2007) and paleontological (Hunt et al. 2008) observations. Because of their rapidity relative to geological timescales, such phenotypic “jumps” are difficult to observe directly (Lande 1985; Kirkpatrick 1982), but their lasting impressions are detectable in both simulated and empirical comparative phenotypic datasets (Figure 1). Among Simpsonian modes of phenotypic macroevolution—including unconstrained gradual evolution, evolutionary stasis, explosive evolution, and punctuated evolution—we found that punctuated evolution is not only detectable, but is a preferred explanation for how body sizes evolved among diverse vertebrate groups. Phenotypic variation accrues in time at a rate that is nearly identical when assuming models of non-stationary gradual evolution versus punctuated evolution; that is, they differ in evolutionary mode rather than evolutionary tempo.

To test for the signatures of different modes of evolution, we assembled neontological comparative datasets for 66 vertebrate clades, composed of time-calibrated phylogenies and body size measurements (Table 1). Importantly, we undertook the call for “broad comprehensive sampling” that is necessary to characterize tempo and mode of evolution without bias (Stanley 1979). We developed a method to fit a model of phylogenetic evolution of continuous traits for processes that model macroevolutionary jumps by exploiting the mathematical properties of Lévy processes. Applying our method to the vertebrate dataset, we found that body size evolution was best explained by jump models in 32% of vertebrate clades when fitting gradual, stationarity, and explosive models of evolution along with four flavors of jump processes. Taking a broader definition of support for rapid evolution, punctuated and Early Burst models are selected for 47% of all clades, and for 72% among those clades with strong model preference of any kind.

Simulated datasets with phylogenetic error do not result in a systematic bias towards preferring Lévy processes over competing models (Figure S2). We found that, if anything, our data filtering and model selection procedures tip the balance to incorrectly favor non-punctuated over punctuated models of evolution. Indeed, the false positive rate seen in simulations are from four to seven times lower than the rate of clades that are best fit by Lévy processes in the empirical data, suggesting a very low false discovery rate. These findings contradicted concerns that gradual and punctuated models would be difficult to distinguish, because Brownian motion models and jump processes result in the same tempo of evolution (as measured by the variance that the processes accumulate per unit time). Nonetheless, rare, large jumps result in heavy-tailed distributions of trait change along branches (as quantified by the excess kurtosis of the trait change distribution), which can be distinguished from gradual evolution occurring at a similar tempo. Turning to the empirical dataset, we found no statistical support that certain model classes were overrepresented within a particular vertebrate group (Table S3). Because each of our five vertebrate datasets was curated under uniform conditions within each group, and because those procedures varied between groups, our empirical results are likely to be robust to ascertainment bias and data quality. Interestingly, this suggests that the five vertebrate groups explore phenotypic space using the same palette of evolutionary modes in roughly equal proportion.

We did not replicate findings of weak support for Early Burst (Harmon et al. 2010b). Several studies have reported that macroevolutionary models of continuous trait evolution may produce misleading results when intraspecific variation is ignored (Cooper et al. 2015; Pennell et al. 2015; Silvestro et al. 2015; Kostikova et al. 2016). Along these lines, we suspected that Early Burst models that ignore intraspecific variation cannot simultaneously explain explosive variation between deeply diverged species in the clade alongside variation concentrated near the tips of the tree. Interestingly, when we forced the model to ignore intraspecific variation, we found that Ornstein-Uhlenbeck to be better supported than Early Burst or Brownian motion (Table S3). This suggests that many findings of support for models of evolutionary stasis over models of adaptive radiations may be due to model misspecification, rather than evidence for adaptive landscapes with single stationary peaks.

We estimate that the median waiting time to jump to new adaptive zones is 10^7.4^ years across clades, consistent with predictions for the time required to jump between adaptive peaks that are separated by fitness valleys via genetic drift in small populations (Lande 1985). Under this model, the primary impediment to peak shifts is escaping the current adaptive zone; once escaped, transitions to new adaptive optima occur rapidly. With biologically reasonable parameters, shifts between adaptive zones are expected to occur every ~10^6^ – 10^7^ generations, but only take ~10^2^ – 10^3^ generations to complete once initiated. Alternatively, this waiting time may represent the timescale of changes of the adaptive landscape itself. Under this model, macroevolutionary jumps are precipitated by changing biotic or abiotic conditions (such as climate change). Standard quantitative genetic theory shows that adaptation to new, nearby optima can occur extremely rapidly, again on the order of ~10^2^ – 10^3^ generations (Kirkpatrick 1982). Future work will be necessary to determine the relative contributions of these two explanations for punctuated evolution to the patterns seen between extant vertebrates.

We found that different kinds of jump processes, which represent different types of pulsed evolution (constant rapid adaptation vs. long periods of stasis broken up by jumps between adaptive zones) leave faint, but unique, signatures in phylogenetic data. By integrating these models into fossil sequences, we suspect that further fine-scale details of macroevolution can be elucidated. Moreover, in quantitative finance, where, for practical purposes, the “fossil record” of stock prices through time is perfectly kept, fine-scale dynamics of jump processes can be inferred (Barndorff-Nielsen et al. 2012; Belomestny 2010; Kappus and Reiß 2010), suggesting that such power exists for suitably densely sampled fossil sequences. However, if rapid evolutionary changes are concentrated near speciation events, as in the classical theory of punctuated equilibrium (Gould and Eldredge 1977), signals for those adaptive shifts may be difficult to locate without a phylogenetic context among fossil sequences (Bokma 2008). Our approach, which uses only modern data but integrates it into a phylogenetic framework, represents an important step towards a fully integrative analysis of macroevolutionary processes.

## 3 Materials and Methods

### 3.1 Likelihoods using characteristic functions

Because most Lévy processes are only known by their characteristic function, we developed a REML algorithm that operates on characteristic functions. We computed the likelihood of the independent contrasts by proceeding from the tips to the root of the phylogeny. To do so, we recursively update the the estimate of the trait at internal nodes as a linear combination of the trait value at the two daughter nodes. We also include an additional term to model the uncertainty in trait values at the tips of the tree due to both intraspecific variation and measurement error. Details of the algorithm can be found in the *SI Appendix.*

### 3.2 Optimizing model fit using REML

To fit the model to data, we developed a package in the R programming language while making key use of several R libraries (Paradis et al. 2004; Bihorel 2015; Pennell et al. 2014). Our package is available at available at https://github.com/Schraiber/punctuatR. Optimal parameter values are obtained by maximizing the likelihood of independent contrasts using the Nelder-Mead simplex method (Nelder and Mead 1965). To ensure that we obtained true maximum likelihood estimates, we performed multiple independent parameter searches (10 for empirical data, 4 for simulated data). In addition, we validated that all maximum likelihood estimates agreed with model hierarchies, e.g. the maximum likelihood for Ornstein-Uhlenbeck must be greater than or equal to that for Brownian motion.

### 3.3 Simulation study for model selection power and sensitivity

We assessed the power and false positive rate of our method by manipulating tree size and phylogenetic error in a controlled simulation setting. Each dataset was simulated by sampling one tree from an empirical posterior distribution of trees (Jetz et al. 2012) then simulating trait data under each of the seven candidate models (BM, OU, EB, JN, NIG, BM+JN, BM+NIG). Simulation parameters are given in Table S1. We then computed AIC weights for each of the seven datasets across the same seven candidate models, once under the originally sampled tree (the “true” tree), then again under the maximum clade credibility tree that summarizes the posterior (the “consensus” tree). Repeating this procedure 100 times, we counted how often each model type is selected in the presence and absence of tree error. To quantify how tree size influences bias in model selection, this experiment was conducted for two clades with sizes representative of our empirical datasets (Procellariidae with 106 taxa; Tityridae, Tyrannidae, and allies with 325 taxa). In total, we simulated 1,400 datasets and performed 39,200 maximum likelihood fittings.

### 3.4 Vertebrate dataset

Time-calibrated phylogenies and body size measurements were assembled for 66 clades across five major vertebrate groups (Table 1).

We compiled numerous phylogenetic (Harmon et al. 2010b; Cowman and Bellwood 2011; Santini et al. 2013a;b; Sorenson et al. 2013; Bloom and Lovejoy 2014; Roxo et al. 2014; Campanella et al. 2015; Dornburg et al. 2015; Martin and Bonett 2015; Pyron and Wiens 2011; Wright et al. 2015; Jaffe et al. 2011; Jetz et al. 2012; Vos and Mooers 2004; Slater et al. 2010; Nyakatura and Bininda-Emonds 2012; Bibi 2013; Mitchell et al. 2014; Schenk et al. 2013; Shi and Rabosky 2015) and trait measurement (Froese and Pauly 2000; De Lisle and Rowe 2013; Feldman et al. 2016; Dunning Jr 2007; Jones et al. 2009) resources to measure body size evolution. The species relationships within each clade were represented with fixed phylogenies with branch lengths measured in millions of years. Body size data were represented by body length for fish, amphibians, and reptiles; and, mass being a function of volume, the cube root of body mass for mammals and birds. Trait measurements were then log transformed, under the assumption that trait evolution operates on a multiplicative scale (Gingerich 2000). Handling the data in this manner permits the comparison of evolutionary parameters between clades because our body size and tree data share a common scale.

Broadly, we applied the same data assembly strategy to all clades within each of the five vertebrate groups. Clades were generally either extracted from large, taxon-rich phylogenies (amphibians, reptiles, birds), or pooled across independently estimated phylogenies (fish, mammals). Availability and quality of trait data varied between vertebrate group, each requiring special handling. **Fish:** Standard, fork, and total lengths were extracted from Fishbase using RFishbase (Boettiger et al. 2012). For each fish clade, we used an allometric regression to convert all taxa to the most frequently observed length type (Gaygusuz et al. 2006). **Amphibians:** Mean female snout-vent lengths were used for the three Anuran clades (Ranoidea, Hyloidea I, and Hyloidea II). Caudata traits were drawn from (Pennell et al. 2014). **Reptiles:** Total lengths were used for all snake (Serpentes) clades. Maximum straight-line carapace lengths were used for turtles (Chelonia). Snout-vent lengths were used for all remaining reptile clades. **Mammals:** Adult unsexed body mass measurements were used. **Birds:** Adult sex-averaged mean body mass measurements were used.

We corrected for two sources of measurement error that would potentially mislead model selection tests to favor Lévy processes. First, rounding error in the trait data caused some clades to contain contrasts that have the exact value of zero. Zero-valued contrasts artificially favor the JN process, which contains a singularity at 0. Second, grossly inaccurate measurements and/or phylogenetic error will drive some portion of contrasts to appear excessively large, fattening the tail density of the trait distribution. Such inaccurate contrasts might result from errors in the trait data or the phylogenetic tree. To mitigate such errors, we first pruned away any subclades with zero-valued contrasts for all clades. Then, assuming a Brownian motion model, we pruned away the subclade with the most extreme valued contrast for 63 of 66 clades. The three clades exempted from the second filtering step had either a large, but expected, contrast (Anguimorpha, involving the contrast containing *Varanus komodoensis*) or had the largest contrast fall near the base of the tree (Procellariidae and Thamnophilidae). The filtering procedures resulted in a 4.7% loss of data (8,323 of 8,729 taxa remained). *SI Appendix* contains further details regarding how the input data were processed.

### 3.5 Empirical model selection

We computed AIC weights (wAIC) for each clade across four model classes: BM, OU, EB, and Lévy processes (Figure 1). The Lévy process with the highest AIC score was chosen to represent all Lévy processes within the class. Model selection required the model class to be at least twice as probable as any competing model class. With Brownian motion being a special case of the remaining six models, this threshold lies on the cusp of the maximum relative probability that it might receive under wAIC. Numerous alternative analyses under various model classifications and assumptions are located in *SI Appendix.*

### 3.6 Expected waiting time between adaptive shifts

For Lévy processes with a jump component, we are interested in the waiting time until a jump outside of the current adaptive zone occurs. Under the symmetric jump models we consider here, the waiting time for a jump larger than *x* is

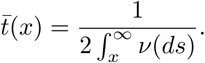

We characterized the width of adaptive zones by the amount of intraspecific variation in a clade. To provide an accurate estimate of intraspecific variation for all clades, we regressed our observed values of *σ*_tip_ against direct measurements of intraspecific standard deviation in birds (Dunning Jr 2007) (n = 16, *p* < 5*e* − 3, *R*^2^ = 0.53, slope = 0.200, intercept = 0.059). Further details for this analysis and results using the uncorrected utip estimates are given in *SI Appendix.*

## 4 Acknowledgments

We would like to thank Damien Wilburn, Tracy Heath, and Ignacio Quintero for helpful comments on an early version of this manuscript. Jonathan Chang, Peter Cowman, and Nathan Upham provided invaluable advice on constructing the empirical datasets. Discussions with Joe Felsenstein helped direct the early course of this project. JGS was supported in the early stages of this work by NSF Postdoctoral Fellowship (DBI-1402120) and subsequently by startup funds from Temple University. MJL was supported initially by the Donnelley Fellowship through the Yale Institute of Biospheric Studies and later through the NSF Postdoctoral Fellowship (DBI-1612153). Analyses were performed on the Yale High Performance Computing clusters.

